# Hair Dermal papilla maintain its hair inducing characteristics via autocrine exosomal miR-23a-3p

**DOI:** 10.1101/2025.07.23.666243

**Authors:** Lei Yan, Jieun Seo, Tatsuto Kageyama, Junji Fukuda

## Abstract

Human dermal papilla cells (hDPCs) are specialized mesenchymal cells that regulate hair follicle development and drive the hair cycle. However, when cultured in vitro, hDPCs progressively lose their hair-inductive capacity. This decline may be closely associated with the absence of the native microenvironmental niche, which plays a critical role in maintaining hDPC identity and function in vivo. In addition to extrinsic cues, autocrine signaling within the dermal papilla itself may serve as a key mechanism for sustaining its regenerative potential. In this study, we present a novel growth model that enables real-time tracking of hDPC functional recovery, with particular attention to autocrine signaling and niche reprogramming. Our results reveal that intrinsic feedback loops are essential for maintaining hDPC function during the anagen phase. By elucidating the dynamic interplay between hDPC expansion and hair-inductive capacity, this work advances our understanding of hair follicle biology and highlights new avenues for developing regenerative therapies for hair loss.

## Introduction

Hair follicle development and cyclic regeneration (anagen, catagen, and telogen) are tightly regulated by reciprocal interactions through various components in the follicular niche^1^. Among these, dermal papilla (DP) is considered to be essential for sustaining hair cycles^2^. DP is located at the base of hair follicle bulbs and composed of highly specialized mesenchymal cells, DP cells (DPCs), which regulates the key processes through interactions with hair follicle keratinocyte^2^. Transplantation of DPCs, but not dermal fibroblasts, presented a remarkable ability to reprogram non-hair-bearing epidermis into a hair follicle fate, revealing the fate regulation function of DPCs through the epithelial–mesenchymal interactions^3^.

In addition to sending signals, DPCs actively receive regulatory cues from a variety of neighboring niche components, including intradermal adipose tissue^4^, blood vessels^5^, immune cells^6^, lymphatic vessels^7^, and nerves^8^ to engage in hair follicle development and cycling. For instance, perifollicular dermal white adipose tissue promotes hair growth by secreting hepatocyte growth factor (HGF), which enhances Wnt/β-catenin signaling in the dermal papilla. Endothelial-derived Apelin-Aplnr signaling, which stimulated DPC proliferation and angiogenesis. Macrophages contribute through extracellular vesicles enriched in Wnt3a/Wnt7b, which activate β-catenin signaling in DPCs, increase LEF1 and VCAN expression, and enhance DPC proliferation and migration, accelerating anagen onset. Similarly, lymphatic vessels and adrenergic nerves influence follicular dynamics via fluid homeostasis and neurotrophic factors, respectively.

While previous studies have primarily focused on interactions between DPCs and their surrounding niche, the mechanisms by which DPCs coordinate synchronized behavior through interactions between cells in DP to regulate hair follicle development and cycling remain largely unexplored. Growth factors secreted by DPCs, such as FGFs and IGFs, contribute to this synchronization through autocrine and paracrine signaling^9^. Recent research has further highlighted the role of DPC-derived exosomes as critical mediators of these signaling. Notably, Hu et al. demonstrated that exosomes from DPC spheroids, enriched in miR-218-5p, significantly promote hair regeneration by downregulating SFRP2 and activating β-catenin signaling^10^. These findings raise the possibility that DPCs may affect their behavior via exosome-mediated signaling, forming a coordinated signaling network essential for effective hair follicle regeneration.

To date, most of these findings have been limited to murine models, and molecular insights into human DPCs remain scarce. A simplified in vitro culture system that more accurately recapitulates the complex in vivo interactions is needed to better understand and mimic human DPC function. The current gold standard—3D spheroid culture—can partially replicate the native morphology and restore some functional properties of hDPCs. However, it falls short of fully mimicking the physiological dermal papilla niche. For example, essential genes for DPCs such as IGF2 and DIO2 are not fully expressed.^11^. Therefore, there is a need for another culture system that better replicates both the molecular identity and regenerative function of hDPCs.

In this study, we discovered that when hDPCs are cultured long-term without trypsinization, they spontaneously self-organize into multilayered structures with heterogenous topographies of cell layers. These structures exhibited a higher expression of hair-inductive genes compared to conventional spheroid cultures. To understand the underlying mechanism, we focused on exosome-mediated autocrine signaling. This culture approach was designed to address how hDPCs maintain their functional identity and regenerative potential over extended periods, both in vitro and potentially in vivo.

## Results

### Characterization of hDPCs in dense-layered culture

In conventional 2D culture, DPCs proliferate rapidly, typically reaching 70–80% confluence within 5 days after seeding. Under these conditions, the cell number increases approximately 7- to 8-fold per passage, and after two passages, the total cell number can expand up to ∼60-fold. However, repeated passaging leads to a progressive decline in DP cell functionality, as indicated by a marked reduction in alkaline phosphatase (ALP) expression (Supplementary Fig. 1). To enhance or maintain DP function, spheroid culture without passaging has been employed. This method preserves or even increases ALP expression, but significantly suppresses cell proliferation.

To improve the physiological relevance of in vitro dermal papilla models, we unintentionally extended the culture period of hDPCs and observed a notable recovery of their functional properties. This led us to establish a dense-layer culture (DLC) system (Fig. 1a). As an initial evaluation, passage 5 hDPCs were cultured in 24-well flat-bottom plates for 30 days without passaging. By day 5, hDPCs reached confluence and continued to proliferate beyond contact inhibition, forming multilayered, cell-dense structures (Fig. 1b). Giemsa staining revealed the emergence of spatial heterogeneity—ridge (5-6 layers) and valley (3-4 layers) regions (Fig. 1d).

**Fig. 1.**
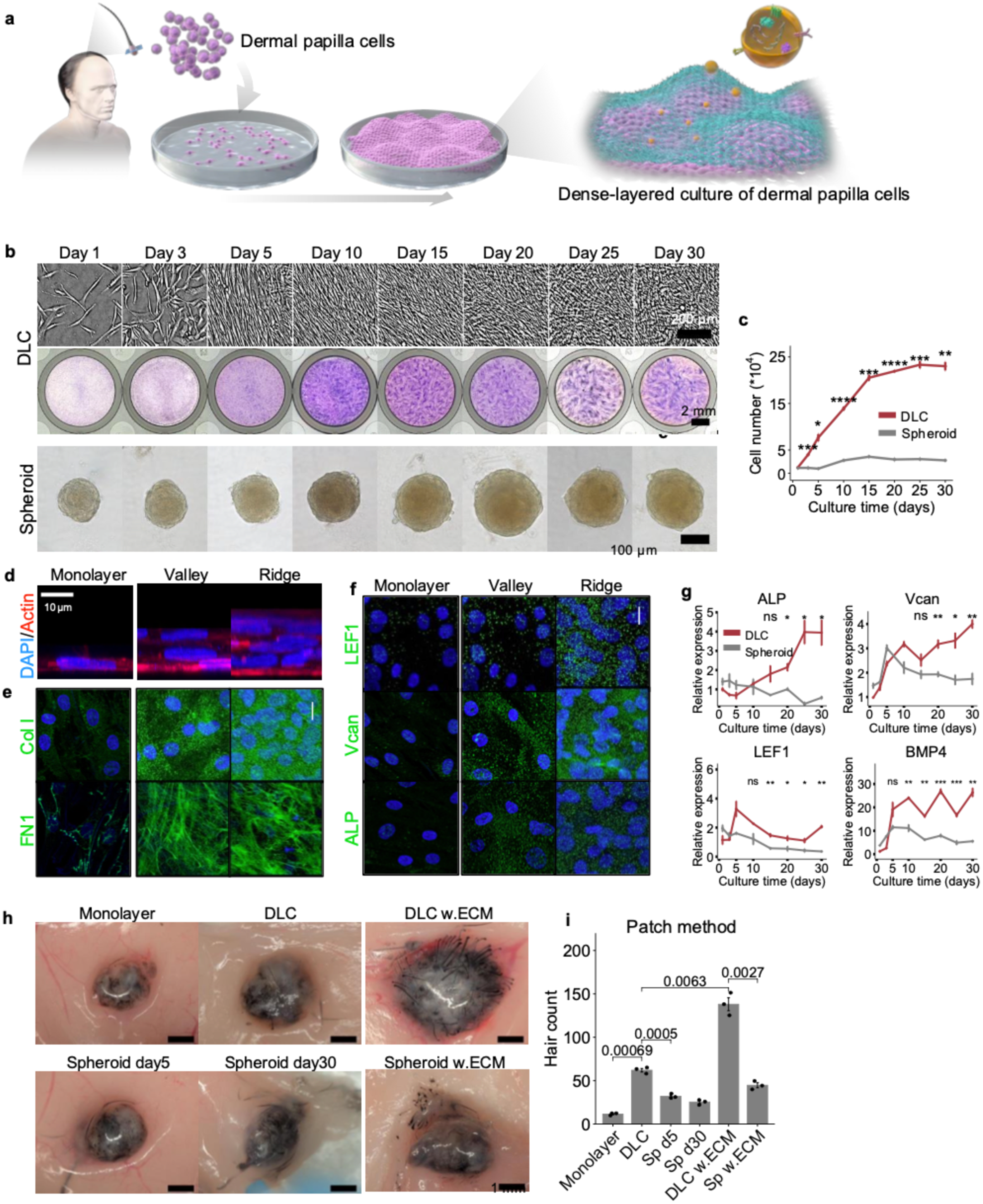
Dense-layered culture of purchased dermal papilla cells. **a** Schematic diagram of densely layered culture of human dermal papilla cells. **b** Morphological transformation of commercially available DLC-treated hDPC and spheroid-cultured hDPC. **c** Cell proliferation rate of DLC- and spheroid-cultured hDPC (n = 3). **d** Structure of monolayer hDPC and heterogeneous valley and ridge structures of DLC-ed hDPC. **e** Immunostaining of DLC-treated hDPC ECM excretion of Fibronectin and Collagen Type I. **f** Immunostaining of ALP, Vcan and LEF expression during DLC. **g** Relative gene expression in DLC- and spheroid-cultured hDPC. **h** Number of hairs generated in the patch assay using fresh hDPC collected from donors under various conditions. The scale bars are indicated in each figure. * P<0.05 ** P < 0.01 ***P < 0.001 ****P < 0.0001 Data represent mean ± SEM of biological replicates (n = 3 in c, d, h). Two-tailed Student’s t-tests (c, d, h).

Compared to spheroid culture, DLC promoted significantly greater cell proliferation, reaching a 23-fold increase by day 30, while spheroids plateaued below 3-fold (Fig. 1c). Time-lapse imaging confirmed dynamic cell movement and accumulation into stable multilayered zones (Supplementary Video 1). Immunostaining demonstrated substantial deposition of extracellular matrix components—fibronectin and collagen I—by day 30 (Fig. 1e). These ECM proteins filled the intercellular spaces and served as a scaffold that supported vertical stacking of cells, contributing to the formation of the multilayered structure observed in DLC cultures.

We next assessed trichogenic marker expression. Immunostaining revealed minimal expression of ALP, Versican (Vcan), and lymphoid enhancer-binding Factor 1 (LEF1) in early monolayers, but robust expression in day 30 DLC (Fig. 1f). QPCR analysis confirmed progressive upregulation of trichogenic genes in DLC, including ALP, Vcan, LEF1, and Bone morphogenetic protein 4 (BMP4), with DLC outperforming spheroid culture by day 30 (Fig. 1g).

In earlier experiments, we used hDPCs that had undergone two rounds of cryopreservation and were expanded up to passage 5, conditions known to significantly reduce their functional capacity. Therefore, the robust recovery observed under DLC conditions may have been influenced, in part, by the low baseline functionality of the starting cells. We also assessed hDPCs that were freshly isolated and pre-cultured without cryopreservation. These cells also showed functional recovery under DLC conditions (Supplementary Fig. 3d), indicating that the beneficial effects of DLC are not limited to cryopreserved cells. To examine the hair-inductive capacity of DLC-cultured hDPCs, we performed patch assays in immunodeficient mice using primary hDPCs derived from donors (Fig. 1h), indicating that the DLC effect is not limited to cryopreserved cells. Notably, transplanting intact DLC sheets with their native ECM— without enzymatic dissociation—further enhanced hair regeneration compared to dissociated DLC. The presence of ECM also appeared to improve hair shaft thickness (Supplementary Fig. 3e). In contrast, ECM retention had minimal effect in the spheroid group, underscoring the functional contribution of DLC-derived ECM.

These results demonstrate that the DLC method not only enhances hDPC proliferation and restores trichogenic gene expression, but also supports robust hair follicle neogenesis in vivo. The role of ECM appears central, enhancing both the quantity and thickness of regenerated hair.

### Elucidation of genes expression recovered in DLC vs spheroid culture

Given that hDPCs cultured using the DLC method exhibited superior hair-inductive potential compared to spheroid culture, we sought to investigate the underlying molecular mechanisms driving this enhancement. We performed gene expression profiling via microarray analysis across six distinct cell conditions: fresh dermal papillae, primarily cultured dermal papillae (Pre), monolayer-cultured (Monolayer) and DLC-hDPC (DLC), and spheroid-cultured hDPC at both day 5 (Spheroid D5), and day 30 (Spheroid D30) (Fig. 2a), derived from three donors (aged 19, 27, and 46). Microarray data were normalized and filtered following the manufacturer’s protocol, resulting in 15,625 genes for downstream analysis. Dimensionality reduction using t-SNE was performed (Fig. 2b). We focused comparisons on two pairs: Fresh vs. DLC and Fresh vs. Spheroid D5.

**Fig. 2.**
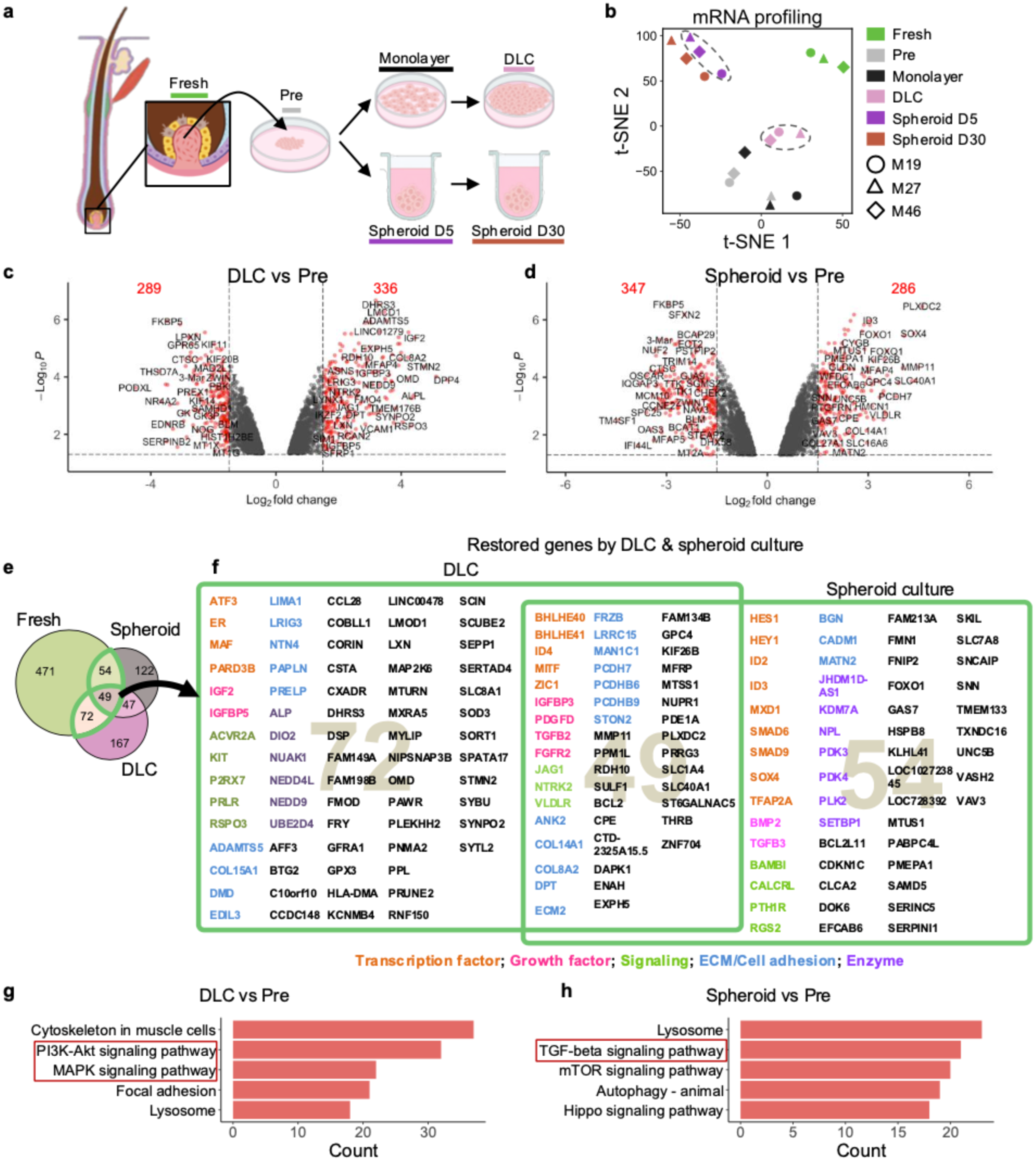
Gene expression analysis of DLC-treated hDPCs and spheroid cultured hDPCs. **a** Schematic illustrating the six experimental conditions used for hDPC microarray analysis. **b** t-SNE plot depicting mRNA expression profiles of three biological replicates for each condition. **c** Volcano plot comparing DEGs between DLC and Pre; **d** Volcano plot comparing DEGs between spheroid culture on day 5 and Pre. **e** Venn diagram showing the overlap between downregulated genes induced by primary culture and genes recovered by DLC and spheroid culture. **f** Expanded view of the genes highlighted in the green box in the Venn diagram. **g, h** KEGG pathway analysis of differentially expressed genes in DLC (g) and spheroid culture (h) compared to Pre.

Volcano plot analysis identified the most significantly differentially expressed genes in both DLC and spheroid cultures compared with Pre (Fig. 2c, 2d). Among the 646 genes downregulated during the primary culture period (Fig. 2e), 121 genes were recovered in the DLC, 49 genes are commonly recovered by DLC and spheroid culture (Fig. 2f).

KEGG (Kyoto Encyclopedia of Genes and Genomes) analysis revealed enrichment of PI3K-Akt and MAPK signaling pathways in DLC hDPCs, suggesting their involvement in enhancing the trichogenic potential of these cells. This finding is consistent with previous reports showing that PI3K-Akt signaling is activated by collagen type I and contributes to the hair-inductive capacity of hDPCs^12^. In the DLC method, the self-excreted extracellular matrix may be responsible for activating this signaling pathway. Enrichment of MAPK signaling further supports its role in maintaining or enhancing DPC functionality^16^. Spheroid culture showed enrichment of TGF-β signaling, which corroborates previous reports^13^.

### Potential pathway for function recovery in DLC

We performed GSEA using the hallmark pathway database on fresh vs. Pre, DLC vs. Pre, Spheroid vs. Pre and DLC vs. spheroid to choose and understand the possible underlying principles of DLC (Fig. 3a). This analysis aimed to identify key signaling pathways and biological processes associated with the maintenance or recovery of trichogenic potential in hDPCs.

**Fig. 3.**
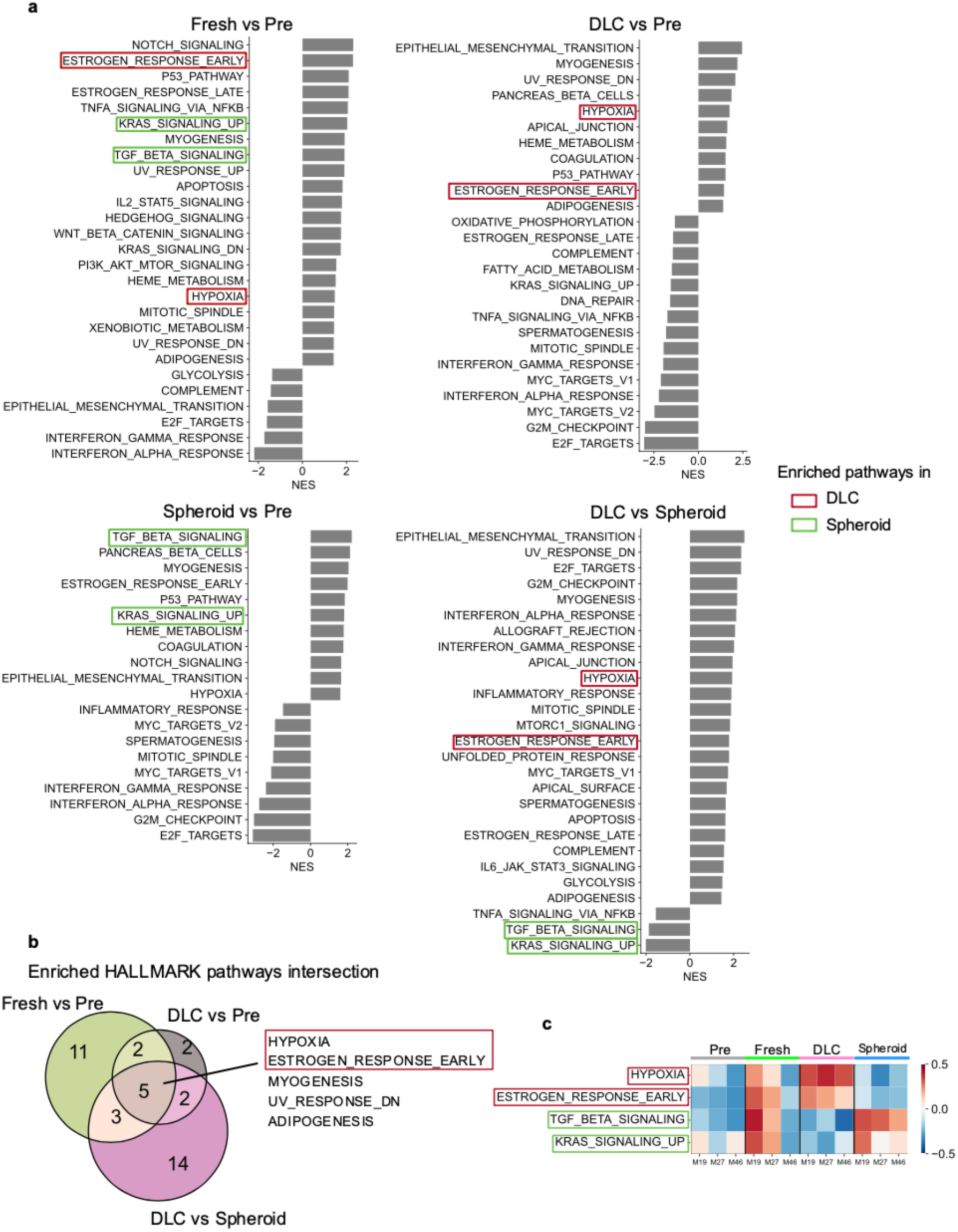
Identification of potential molecular mechanisms underlying DLC. **a** GSEA results comparing Fresh vs Pre, DLC vs Pre, Spheroid vs Pre, and DLC vs. Spheroid conditions, highlighting differentially activated pathways potentially responsible for DLC’s mechanism **b** Venn diagram showing the intersection of five possible molecular mechanisms identified for DLC **c** GSVA results for each patient, visualizing the activity of signaling pathways of interests.

The intersection of the gene sets differentially expressed between fresh vs. Pre, DLC vs. Pre, and DLC vs. spheroid cultures highlighted two key pathways: hypoxia and early estrogen response (Fig. 3b). Hypoxic signaling has previously been shown to enhance the hair inductivity of hDPCs^14^. Estrogen treatment has been demonstrated to promote hair regrowth in and in animal models^15^ and AGA patients^16^. The elevated hypoxia-related signaling observed in DLC cultures suggests that the dense, multilayered structure may create a localized hypoxic microenvironment that activates this pathway. The mechanism by which estrogen signaling is induced in DLC will be further explored in the next section.

While GSEA provides an average enrichment score across samples, we also applied Gene Set Variation Analysis (GSVA) to assess pathway activity at the individual sample level. specifically, the GSVA showed that the score for the 19-year-old male sample was the highest among the three, demonstrating a gradient from the ages of 19 to 27 and 46 (Fig. 3c). Despite these variations in the fresh samples, hDPCs cultured under DLC conditions consistently showed increased activation of hypoxia pathways across all donor ages. This suggests that the DLC method exerts a robust, age-independent effect on the activation of signaling pathways associated with trichogenic potential.

### Hypoxia signaling and estrogen signaling of DLC

To further confirm the activation of hypoxia and estrogen signaling during DLC culture, we performed quantitative PCR analyses. The results demonstrated a significant upregulation of HIF1α and estrogen receptor (ER) expression in DLC-cultured hDPCs compared to spheroid cultures (Fig. 4a, 4b). HIF1α expression began to rise significantly after day 10, coinciding with the formation of a multilayered cell structure, suggesting that local hypoxia may result from increased cell density in DLC.

**Fig. 4.**
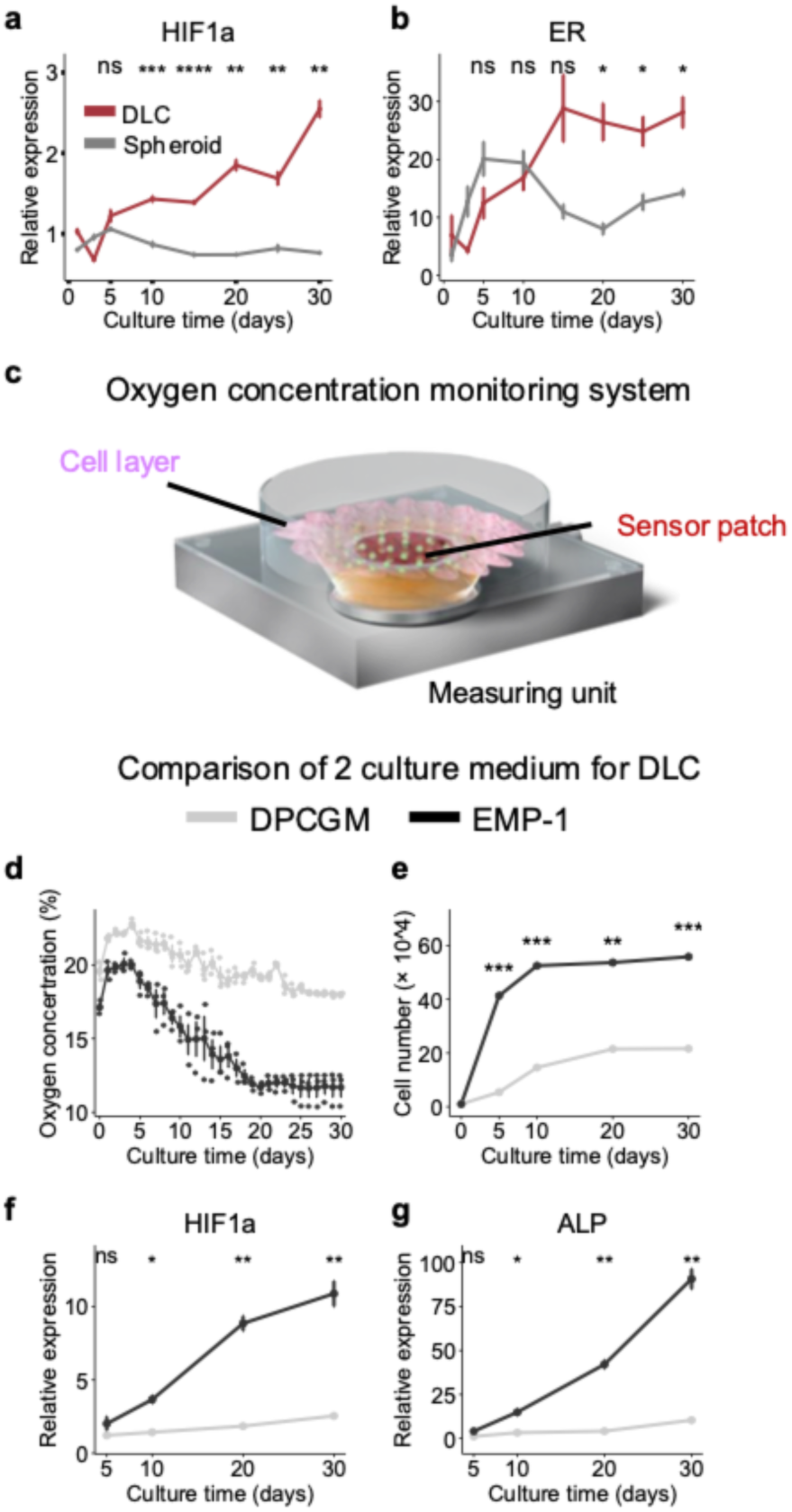
Validation of potential candidates in DLC culture. **a, b** Time-series analysis of HIF-1α and ERα expression levels in human dermal papilla cells (hDPCs) cultured under DLC and spheroid conditions. **c** Schematic representation of the oxygen sensor (THA-450-16SL) used to monitor oxygen levels in the culture system. (i) Comparison of oxygen concentrations in the culture media between DPCGM and EMP-1 over time. (ii) The proliferation dynamics of hDPCs in DPCGM versus EMP-1 are presented as cell numbers over the culture period. (iii, iv) Relative expression levels of HIF-1α (iii) and ALP (iv) in hDPCs cultured with DPCGM and EMP-1. Statistical significance was denoted as ns, not significant; *p < 0.05; **p < 0.01; ***p < 0.001.

**Fig. 5.**
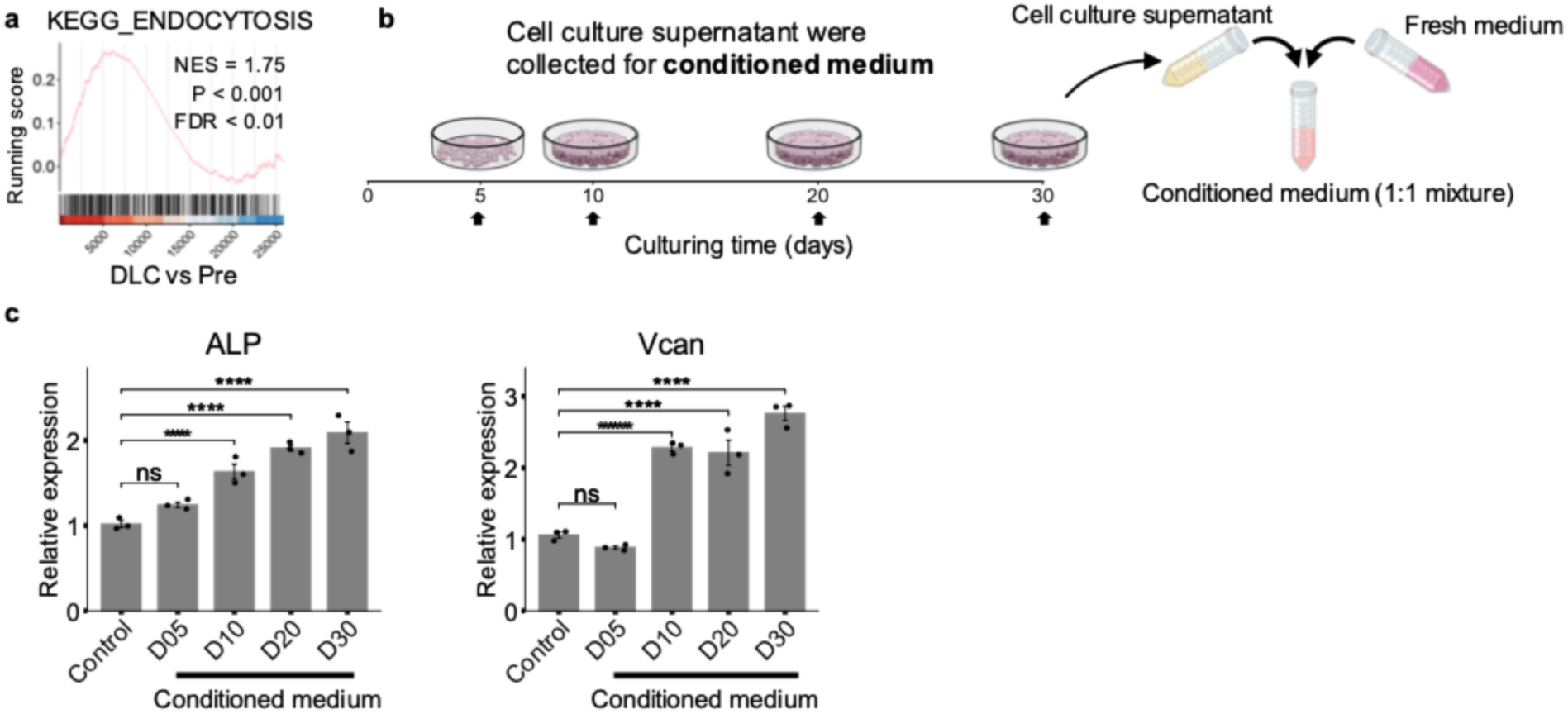
Analysis the supernatants’ effect derived by DLC treated hDPC. **a** DLC-treated hDPCs revealed enrichment of the KEGG_ENDOCYTOSIS pathway. **b** Schematic illustration of the composition of conditioned medium collected from DLC-treated hDPCs. **c** ALP and Vcan expression in recipient cells was enhanced by treatment with conditioned medium derived from DLC-treated hDPCs.

To enhance hypoxia resulting from cell stacking in dense-layer cultures, we tested an alternative culture medium, EMP-1, which is known to promote cell proliferation. During the 30-day DLC period, hDPCs cultured in EMP-1 expanded approximately 40-fold by day 5 and over 50-fold by day 10, whereas cells cultured in DPCGM showed a 21-fold increase by the end of the culture period (Fig 4e). To assess the oxygen partial pressure in each culture condition, we employed a highly sensitive optical fiber oxygen sensor incorporating a Pt(II) complex embedded in a sol-gel matrix^17^ (Fig. 4c). Sensors were immobilized in 24-well plates, and hDPCs were cultured on top for 30 days.

Measurements revealed a ∼10% reduction in oxygen concentration with DPCGM, compared to an approximately 50% reduction under EMP-1 conditions. In contrast to DPCGM, the expression level of HIF-1α exhibited an over 10-fold increase compared to monolayer culture, and DPCGM showed a three-fold increase. While the expression level of ALP increased five-fold using DPCGM, EMP-1demonstrated an surprising over 90-fold upregulation of ALP expression. These findings underscore the influence of hypoxia and the efficacy of EMP-1 in inducing substantial changes in oxygen levels and gene expression.

### The effect of supernatant conditioned medium

In the comparison of microarray gene profiles, we noted significant upregulation of scinderin (SCIN) and NEDD4 like E3 ubiquitin protein ligase (NEDD4L) in response to DLC treatment (Fig. 2f). SCIN, initially identified in chromaffin cells of the bovine adrenal medulla, acts as an actin-depolymerizing agent that disassembles the cortical network of actin filaments in response to bursts of intracellular calcium, leading to the exocytosis of secretory vesicles^18^. NEDD4L, known for its involvement in endocytosis^19^, also exhibited increased expression, highlighting its potential correlation with DLC-induced changes. GSEA based on the KEGG pathway for endocytosis confirmed the involvement of endocytic processes in DLC-treated hDPC, suggesting the possibility of autocrine signaling. To assess the impact of the secretome, we collected supernatants at different culture time points (5, 10, 20, and 30 days) and mixed them with fresh culture medium at a 1:1 ratio. Subsequent cell culture for 3 days revealed a noteworthy upregulation in gene expression, particularly for ALP (1.63-, 1.91-, and 2.09-fold upregulation) and Vcan (2.28-, 2.21-, and 2.76-fold upregulation) in the supernatants from days 10, 20, and 30, respectively. These findings strongly indicate an autocrine effect mediated by factors secreted by hDPCs during DLC.

### Analysis the component and effect of exosome released by dense-layer cultured DPC

Given the close association between endocytosis and exosomes, we collected and characterized exosomes from DLC-treated hDPCs at different time points (5, 10, 20, and 30 days) (Fig. 6a). We observed a general trend of decreasing exosome diameters and increasing uniformity over time (Fig. 6b). Exosomes are crucial mediators of intercellular communication that facilitate the transfer of microRNAs (miRNAs) and other bioactive molecules between cells. MiRNAs are potent regulators of gene expression and play essential roles in various cellular processes, including the differentiation and proliferation of hDPCs. Accumulating evidence has highlighted the potential of DPC-derived exosomes to promote hair follicle development and growth. To explore the involvement of exosomes in DLC-mediated effects, we collected exosomes from the supernatants of DLC-treated hDPCs and performed miRNA profiling. The T-SNE plot revealed a significant shift in exosome miRNA profiles from days 5 to 30, with days 10, 20, and 30 exhibiting similar profiles (Fig. 6c).

**Fig. 6.**
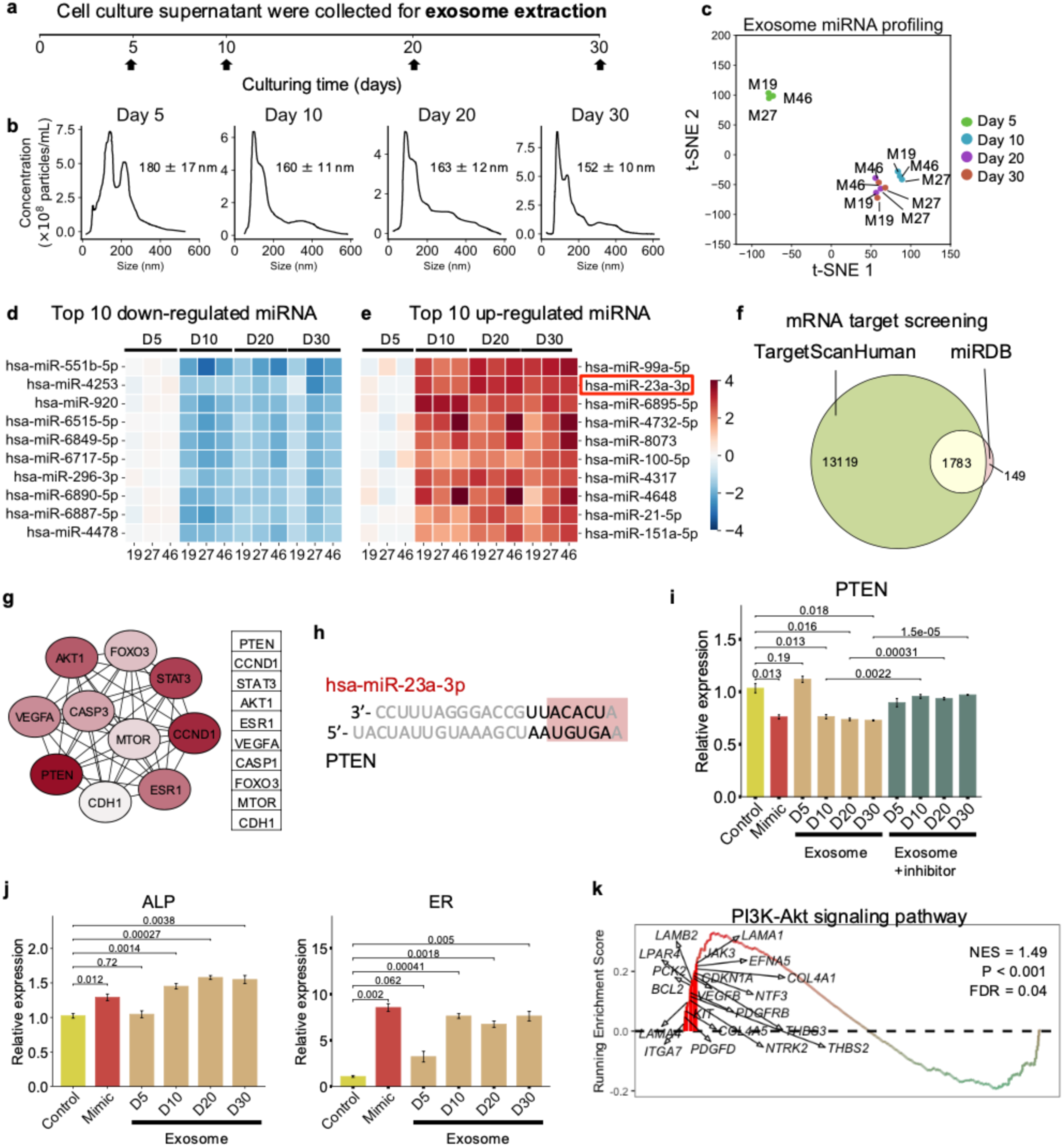
Exploration of DLC-derived exosome. **a** Schematic representation of the DLC-derived exosome extraction process. **b** Exosome size distribution and concentration from culture days 5, 10, 20, and 30. **c** t-SNE profiling of exosomal miRNA, illustrating temporal changes across culture days. **d** Top 10 downregulated miRNAs identified in exosomes at different time points. **e** Top 10 upregulated miRNAs identified in exosomes at different time points. **f** mRNA target screening of the top 10 downregulated and upregulated miRNAs using TargetScanHuman and miRDB. **g** Protein-protein interaction network of top 10 up- and downregulated miRNA target mRNA **h** Interaction of hsa-miR-23a-3p with PTEN **i** Effects of DLC-derived exosomes on PTEN expression in hDPCs, including inhibitor validation. **j** Evaluation of DLC-derived exosomes and hsa-miR-23a-3p’s impact on hDPC ALP and ER expression. **k** GSEA of the PI3K/AKT signaling pathway, revealing the functional secretome of DLC.

Comparative analysis of the top 10 upregulated and top 10 downregulated miRNAs identified on day 30 versus day 5 revealed significant changes in the exosome miRNA profiles (Fig. 6d, 6e, Supplemental Fig. 5). To identify potential target genes, we used the miRNA target prediction databases TargetScan^20^ and miRDB^21^ to identify 1,783 common target genes with a miRNA score exceeding 80 out of 1,932. A protein-protein interaction network was then constructed for these predicted target genes using MCODE, and the most important module was identified based on its high score. This module consisted of 10 nodes and exhibited significant interactions among its genes (Fig. 6g). Further analysis using CytoHubba identified PTEN as the most significant hub gene within this module (Fig. 6g). Among the 20 differentially expressed miRNAs, hsa-miR-23a-3p emerged as a candidate of particular interest.

Our findings demonstrate that the addition of exosomes collected on days 10, 20, and 30 reduced PTEN expression, whereas the miR-23a-3p inhibitor counteracted this effect (Fig. 6i). Furthermore, we established that miR-23a-3p contributes to the restoration of ALP expression and activation of estrogen signaling pathways (Fig. 7i, Fig. 4b).

**Fig. 7.**
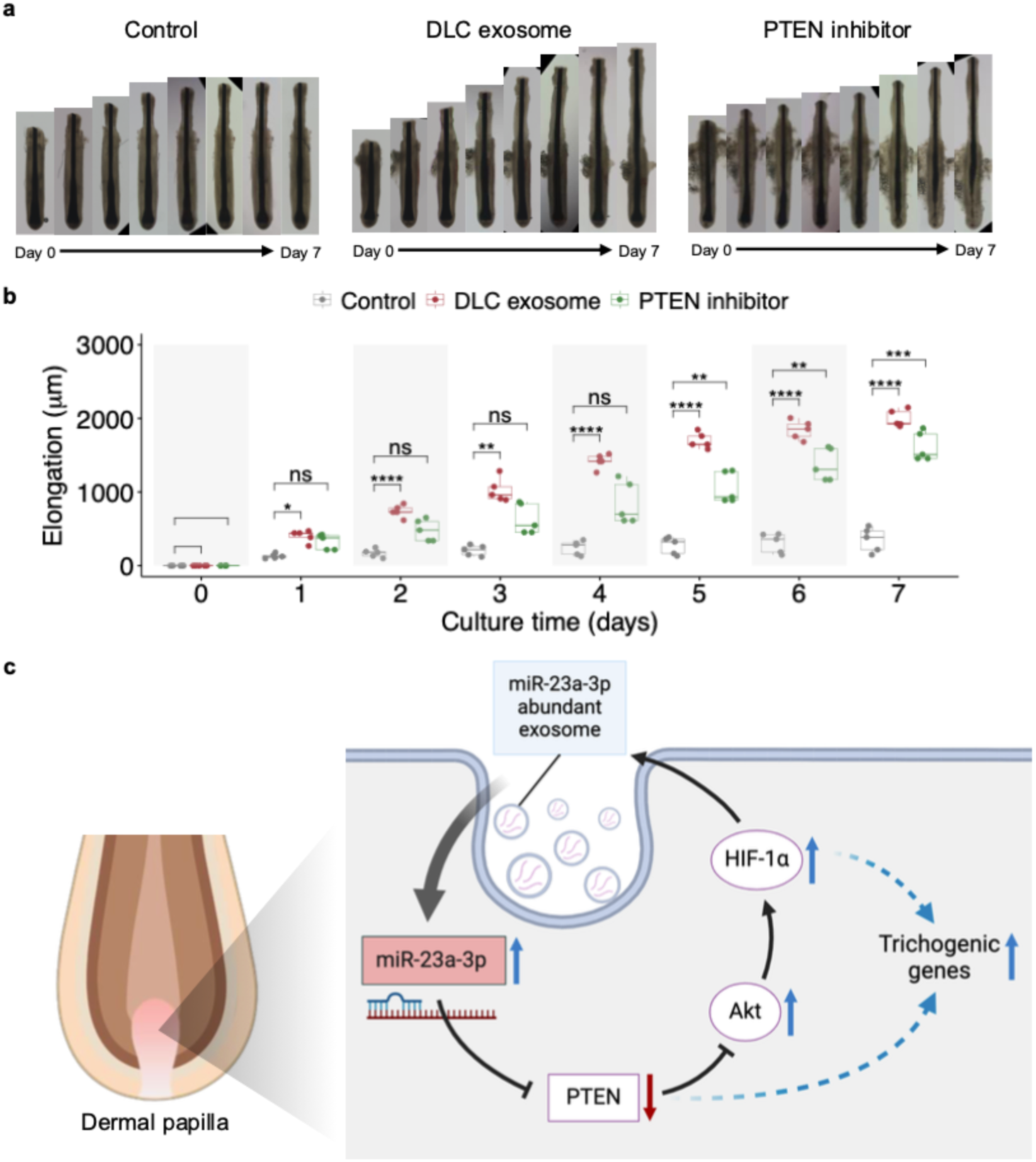
Organ culture and mechanism of DLC. **a, b** Human hair follicles were cultured ex vivo for 7 days to assess the effects of DLC-exo and PTEN inhibition on hair shaft elongation. **c** Schematic illustration of the proposed mechanism by which DLC enhances trichogenic gene expression. In the DLC environment, hypoxia signaling promotes the secretion of exosomal miR-23a-3p by hDPCs. This miRNA contributes to functional restoration by downregulating PTEN, thereby reinforcing hypoxia signaling through a positive feedback loop and supporting the upregulation of trichogenic genes.

PTEN is a key factor in the PI3K/AKT signaling pathway. Our results confirmed that DLC treatment enhanced PI3K/AKT activity (Fig. 6k, Fig. 2g). Although the autocrine effect of DLC likely involves exosome secretion, other growth factors may also be involved. Our analysis revealed the upregulation of growth factors asso6k). However, a more comprehensive secretome analysis is necessary to fully elucidate the factors involved in the autocrine effects of DLC-treated hDPCs. In the present study, we focused on demonstrating the specific effects of exosomal miR-23a-3p.

### DLC-exo’s effect on hair growth

Lastly, we evaluated the effect of DLC-exo on hair growth using a trimmed human hair follicle organ culture model (Fig. 7a). Treatment with DLC-exo significantly promoted hair shaft elongation (Fig. 7b). Additionally, application of the PTEN inhibitor SF1670 further enhanced hair growth, revealing the pivotal role of PTEN signaling in regulating hDPC function and follicular activity. These findings suggest that both DLC-exo treatment and PTEN inhibition represent promising strategies for restoring or enhancing the hair-inductive capacity of hDPCs.

## Discussion

DPCs rapidly lose their hair-inductive functionality once isolated from their native microenvironment. Several strategies have been employed to partially restore their inductive capacity. These include mimicking the in vivo niche using ECM scaffolds such as collagen, activating key signaling pathways (e.g., FGF and Wnt) through the addition of growth factors^22^ or small molecules like fibroblast growth factor or CHIR99021^23^, co-culturing with keratinocytes or using keratinocyte-conditioned medium, and applying physical stimuli such as light or electrical stimulation^24,25^. Among these, 3D spheroid culture has emerged as the current gold standard for maintaining DPC function in vitro^11^.

In this study, we evaluated the effectiveness of the DLC method using both commercially available frozen hDPCs and hDPCs derived from patients with AGA. Remarkably, the DLC method enhanced the functional recovery of both cell types, as demonstrated by genome-wide expression profiling and in vivo hair regeneration assays. Under DLC conditions, hDPCs self-organized into scaffold-free, multilayered cell sheets that secreted abundant ECM components such as fibronectin and collagen type I—both of which are known to support DPC identity and modulate the hair cycle^26^. The self-secreted ECM likely provided the structural and biochemical cues needed to restore native DPC properties. An important feature of the DLC system is the spontaneous formation of a hypoxic microenvironment, without the need for external manipulation. Transcriptomic analysis revealed activation of hypoxia- and estrogen-related signaling pathways during the culture process. Notably, enhancing hDPC proliferation further amplified hypoxic signaling, thereby promoting functional recovery^14,27^. Given the known role of mesenchymal cell sheet cultures in enhancing cytokine secretion^28^, we also investigated the exosomal profile of DLC-cultured hDPCs to better understand the molecular underpinnings of this recovery process.

Mechanistically, we identified a regulatory feedback loop involving HIF-1α, miR-23a-3p, and PTEN. HIF-1α upregulates miR-23a-3p^29^, which in turn suppresses PTEN, thereby sustaining HIF-1α signaling^30^. This feedback loop plays a critical role in maintaining a hypoxic response and enhancing the hair-inductive properties of hDPCs. Moreover, exosomal miR-23a-3p was shown to contribute to early estrogen signaling and may hold therapeutic potential in the context of hair regeneration.

While cell-based therapies for AGA are promising, their clinical translation is frequently hampered by the loss of cell function during in vitro expansion^31^. The DLC method addresses this challenge by enabling the large-scale expansion of hDPCs with restored hair-inductive capacity. Extended culture under DLC conditions promotes spontaneous self-organization, ECM accumulation, and the activation of key signaling pathways—particularly hypoxia-related and early estrogen response pathways.

Furthermore, exosomes isolated from DLC-cultured hDPCs successfully restored hair growth-associated characteristics in recipient cells in vitro. These findings open the door to future studies focused on evaluating the in vivo efficacy of hDPC-derived exosomes as a therapeutic strategy for hair regeneration.

## Methods

### Human dermal papilla cell preparation and DLC method

Commercially available human follicular dermal papilla cells (P2, C-12071, PromoCell) were purchased and subcultured using Follicle Dermal Papilla Cell Growth Medium (DPCGM, C-26502, PromoCell) until passage 4 and were then cryopreserved in cryoprotective freezing medium (Cellbanker 1, Zenoaq, Fukushima, Japan) for later use.

Primary dermal papillae were extracted from patients with androgen alopecia (19, 27, and 46 years of age). All experimental procedures were performed in accordance with the protocols approved by the Institutional Ethical Committee and the YNU Ethical Committee for Medical and Health Research (Authorization No. Hitoi-2018-16), and by following the ethical guidelines for medical and health research involving human subjects from the Ministry of Education, Culture, Sports, Science and Technology and the Ministry of Health, Labour and Welfare, Japan. The patient provided informed consent to collect and use hair follicles damaged during transplantation. Hair follicles were sorted and trimmed under a microscope to remove appendage adipose and epithelial tissues.

Intact dermal papillae were micro-dissected from the follicles and treated with a solution containing 4.8 U/mL dispase II and 100 U/mL collagenase type I for 30 min at 37 °C. The treated dermal papillae were suspended in AmnioMAX-II complete medium and placed in a collagen-coated 6 well plate. After one week of culture, primary hDPCs were harvested by treatment with 0.25% trypsin for 2 min. These were considered precultured hDPC (Pre). Both patient-derived and purchased P5 hDPCs were seeded at 1×10^4^ cells/well in 24-well chamber plate.

DLC methods were performed using Falcon® 24-Well Flat-Bottom Plate (Corning, NY, USA) and spheroid culture was performed using Prime Surface 96U Plate (MS-9096UZ, SUMITOMO BAKELITE, Tokyo, Japan). During day 30 of DLC, 500 µL of DPCGM were exchanged every 2 days. Spheroid culture medium exchange was conducted every two days for half exchange. Both DLC and spheroid culture were conducted at 37 °C under 5% CO2 incubator conditions (MCO-18AIC, SANYO, Osaka, Japan).

### Cell number quantification and heterogeneity visualization of DLC

Cell numbers were quantified using RealTime-Glo™ MT Cell Viability Assay (Madison, WI, USA) following manufacture’s instruction. One-well of the DLC hDPC or 10-wells of spheroids were collected. The luminescence signal was measured using a SpectraMax i3x multi-mode microwell plate reader (Molecular DEVICES).

Giemsa staining was used to visualize the heterogeneous cellular structures. First, the culture medium was discarded, and the cells were rinsed with methanol. The cells were fixed in methanol for 10 min. Giemsa staining solution was prepared by diluting Giemsa azure eosin methylene blue solution (Merck, Burlington, MA, USA) at 1:20 with MilliQ water. After discarding the methanol fixation, 1 mL of Giemsa staining solution was added and incubated for 1 h. The sample was then thoroughly washed under running tap water for 10 min and air-dried for two days.

### Immunostaining

Immunostaining was performed using the following primary antibodies, including Anti-LEF1 antibody (ab137872; Abcam, Cambridge, UK) at a dilution of 1:200, anti-ALP (ab224335; Abcam) at a dilution of 1:200, anti-versican (ab19345; Abcam) at a dilution of 1:200, Anti-Collagen I (ab34710; Abcam) at a dilution of 1:100, and anti-fibronectin (ab2413; Abcam) at a dilution of 1:200. The secondary antibody used was Goat Anti-Rabbit IgG H&L (Alexa Fluor® 488) (ab150077) at a dilution of 1:500. DAPI (D9542; Sigma-Aldrich) was used for nuclear staining. Immunostained samples were visualized under a confocal microscope (Axio Observer Z1 LSM 700; Zeiss, Oberkochen, Germany).

### Gene expression analysis

Total RNA was extracted from samples using a RNeasy mini kit (Qiagen, Netherlands), and cDNA was synthesized via reverse-transcription using a PrimeScript™ RT Master Mix (Perfect Real Time) (Takara-bio, Japan), according to the manufacturer’s instructions. qPCR was performed using the Thermal Cycler Dice Real Time System III (Takara-bio, Japan), with SYBR Premix Ex Taq II (Takara-bio, Japan), and primers for GAPDH (TGGAAATCCCATCACCATCTTC, CGCCCCACTTGATTTTGG), ALP (ATTGACCACGGGCACCAT, CTCCACCGCCTCATGCA), Wnt5a (TCCACCTTCCTCTTCACACTGA, CGTGGCCAGCATCACATC), BMP4 (GCCCGCAGCCTAGCAA, CGGTAAAGATCCCGCATGTAG), HIF-1α (AGTCTGCAACATGGAAGGTATTGC, TCAGCACCAAGCAGGTCATAGG), FN1 (ACAACACCGAGGTGACTGAGAC, GGACACAACGATGCTTCCTGAG), COL1 (GATTCCCTGGACCTAAAGGTGC, AGCCTCTCCATCTTTGCCAGCA), Ki67 (GAAAGAGTGGCAACCTGCCTTC, GCACCAAGTTTTACTACATCTGCC), NOG (CTGGTGGACCTCATCGAACA, CGTCTCGTTCAGATCCTTTTCCT), Vcan (GGCAATCTATTTACCAGGACCTGAT, TGGCACACAGGTGCATACGT), ER (AACCATCACTGAGGTGGCCC, GCTCCAGCTCGTTCCCTTGG), PTEN (TGAGTTCCCTCAGCCGTTACCT, GAGGTTTCCTCTGGTCCTGGTA). All gene expression levels were normalized to those of GAPDH. Relative gene expression was determined using the 2Ct method and presented as the mean ± standard deviation of independent experiments. Statistical evaluation of numerical variables was conducted using Student’s t-tests, and differences with p values less than 0.05 were considered statistically significant.

### Patch assay hair induction transplantation

Pregnant C57BL/6 mice (CLEA Japan, Inc.) were used to obtain embryonic mouse dorsal epithelial cells. For host transplantation, ICR nude mice were purchased from Charles River Laboratories, Japan. The researcher attended Animal Ethics Committee training, and all experiments adhered to institutional animal welfare regulations. Mice were maintained under pathogen-free conditions for 21 days following transplantation.

To prepare for transplantation, the mice were anesthetized, and the dorsal skin was sanitized with tincture iodine and 70% ethanol. Shallow stab wounds were created on the dorsal skin using a 20-G ophthalmic V-lance (Alcon, Tokyo, Japan). A mixture of 2×10⁵ human dermal papilla cells (hDPCs) and 2×10⁵ embryonic mouse dorsal epithelial cells was transplanted into the prepared sites.

Post-transplantation, the generated hair cuticles were examined using a scanning electron microscope (TM-1000, Hitachi, Tokyo, Japan) at an accelerating voltage of 5–10 kV. To quantify the number of hairs, clumps were enzymatically dissociated using a 100 U/mL collagenase solution, and individual hairs were manually counted under a stereomicroscope (Mantis, Vision Engineering, Woking, UK).

### Microarray profiling and statistical analysis

For microarray analysis, total RNA was isolated from fresh and cultured hDPC using an RNeasy Mini Kit (Qiagen, Netherlands) following the manufacturer’s instructions.

Amplification of RNA was performed using GeneChip™ 3’ IVT Pico Kit (Catalog number: 902789, Applied Biosystems™) according to the manufacturer’s protocol. The amplified RNA was hybridized to a Human Genome U133 Plus 2.0 Array (Thermo Fisher Scientific). In all the experiments, initial data preprocessing was conducted by Kurabo Industries Ltd. (Japan), and the data were displayed using Transcriptome Viewer (Kurabo, Tokyo, Japan).

### Bioinformatics analysis

Bioinformatics analysis was conducted using R software (version 4.2.3). Differentially expressed genes were analyzed using the limma package^32^ (version 3.54.2). GSEA and KEGG enrichment analyses were performed using the ClusterProfiler package^33^ (version 4.6.2). The GSVA was performed using the GSVA package^34^ (version 1.46.0).

### Oxygen concertation measurement

Oxygen concentration was measured using the THA-450-16SL unit and sensors provided by Able Biott JAPAN Co., Ltd (Tokyo, Japan). The measuring unit was calibrated prior to use. Calibration for 0% oxygen concentration was performed using Milli-Q water mixed with 2% sodium dithionite, while Milli-Q water alone was used to calibrate for an oxygen concentration of 22%.

### Cell culture supernatant, exosome extraction, nanoparticle tracking analysis and miRNA profiling analysis

Cell culture supernatants were collected from DPCGM on days 5, 10, 20, and 30, following a 48-hour exposure to the cells. To prepare the conditioned medium, the collected supernatants were mixed with fresh culture medium at a 1:1 ratio. The effect of the conditioned medium was assessed by culturing cells in it for 3 days.

For exosome extraction, DPCGM containing FBS was replaced with an FBS-free culture medium, EMP-1 (Rohto, Osaka, Japan). The effect of DLC was initially demonstrated using hDPCs cultured in EMP-1 medium. Supernatants from the cultured hDPCs were collected and centrifuged at 5000 rpm for 30 minutes. Exosomes were then isolated using the Total Exosome Isolation Reagent (from cell culture media) (Invitrogen, Carlsbad, CA, USA) according to the manufacturer’s protocol. The isolated exosomes were characterized using transmission electron microscopy (TEM) and nanoparticle tracking analysis (NanoSight NS300; Malvern Panalytical, Worcestershire, UK).

Microarray analysis of the exosomes was conducted using 3D-gene human miRNA oligo chips (ver.21; Toray Industries, Tokyo, Japan) to investigate the expression of miRNAs in the exosomes.

### miRNA transfection

For the miRNA transfection experiments, we utilized Lipofectamine™ RNAiMAX Transfection Reagent (Thermo Fischer, MA, USA) and Opti-MEM™ I Reduced Serum Medium (Gibco). The mimic for hsa-mir-23a-3p was MC10644 (Thermo Fisher Scientific, Waltham, MA, USA) and the inhibitor was MIMAT0029481 (Thermo Fisher Scientific). All procedures were performed according to manufacturer’s instructions.

### Hair follicle organ culture

Human scalp hair follicles were obtained from patients with androgenic alopecia after informed consent was obtained. This study was approved by the ethics committees of the Kanagawa Institute of Industrial Science and Technology (approval number: S-2019-01) and Yokohama National University (approval number: 2021-04), and was conducted in accordance with institutional guidelines and the Declaration of Helsinki. Scalp hair follicles were isolated from androgenic alopecia (AGA) patients and cultured in low-adhesion 24-well plate. The culture medium consisted of Advanced DMEM/F-12 (Thermo Fisher Scientific, Waltham, MA, USA) supplemented with 1% GlutaMAX-I (Thermo Fisher Scientific) 0.2% Normocin (InvivoGen, San Diego, CA, USA) and 1% Penicillin-Streptomycin (10,000 U/mL, Gibco™, Waltham, MA, USA). Experimental groups included: (1) a control group with no additional treatment, (2) a DLC-exosome (DLC-exo) group treated with 20 µg/mL total protein concentration, and (3) a PTEN inhibitor group treated with 2 µM SF1670 (Selleck, Houston, TX, USA). Hair shaft elongation was measured using ImageJ software.

### Statistical analysis

All numerical values represent mean ± Standard error of the mean unless stated otherwise. Statistical significance was determined using two-tailed Student’s t-test and one-way ANOVA with Tukey’s multiple comparison test. For transcriptome analysis, Benjamin–Hochberg multiple comparison adjustment was applied, and the corrected p-value <0.05 was considered statistically significant.

## Acknowledgements

This work was partially supported by the Kanagawa Prefectural Institute of Advanced Industrial Science and Technology (KISTEC), the Japan Society for the Promotion of Science (JSPS) Kakenhi grants (project numbers 23K13615 and 23K26464), and the Japan Agency for Medical Research and Development (AMED) (grant number 23bm1123031h0001).

We thank Dr. Yinghui Zhou, Professor Yusuke Sato (Tohoku University, Japan), and Professor Atsushi Suzuki (Yokohama National University, Japan) for their support in animal and exosome experiments; Dr. Binbin Zhang Molino and Dr. Paul Molino for appliance support; Dr. Keiichiro Kasai and the clinical staff at the Shonan Beauty Clinic (SBC) Shinjuku institution for facilitating this research by offering human hair shaft samples.

## Author contributions

L.Y. conducted the conceptualization, experiments, data visualization, and manuscript writing. T.K. contributed to manuscript writing and provided technical support for experiments. J.S. participated in both writing and experimental work. J.F. contributed to the conceptualization, supervised the project, performed fact-checking.

## Additional information

### Data availability

All raw microarray data will be made publicly available through the NCBI database.

---

**Supplementary fig. 1.**
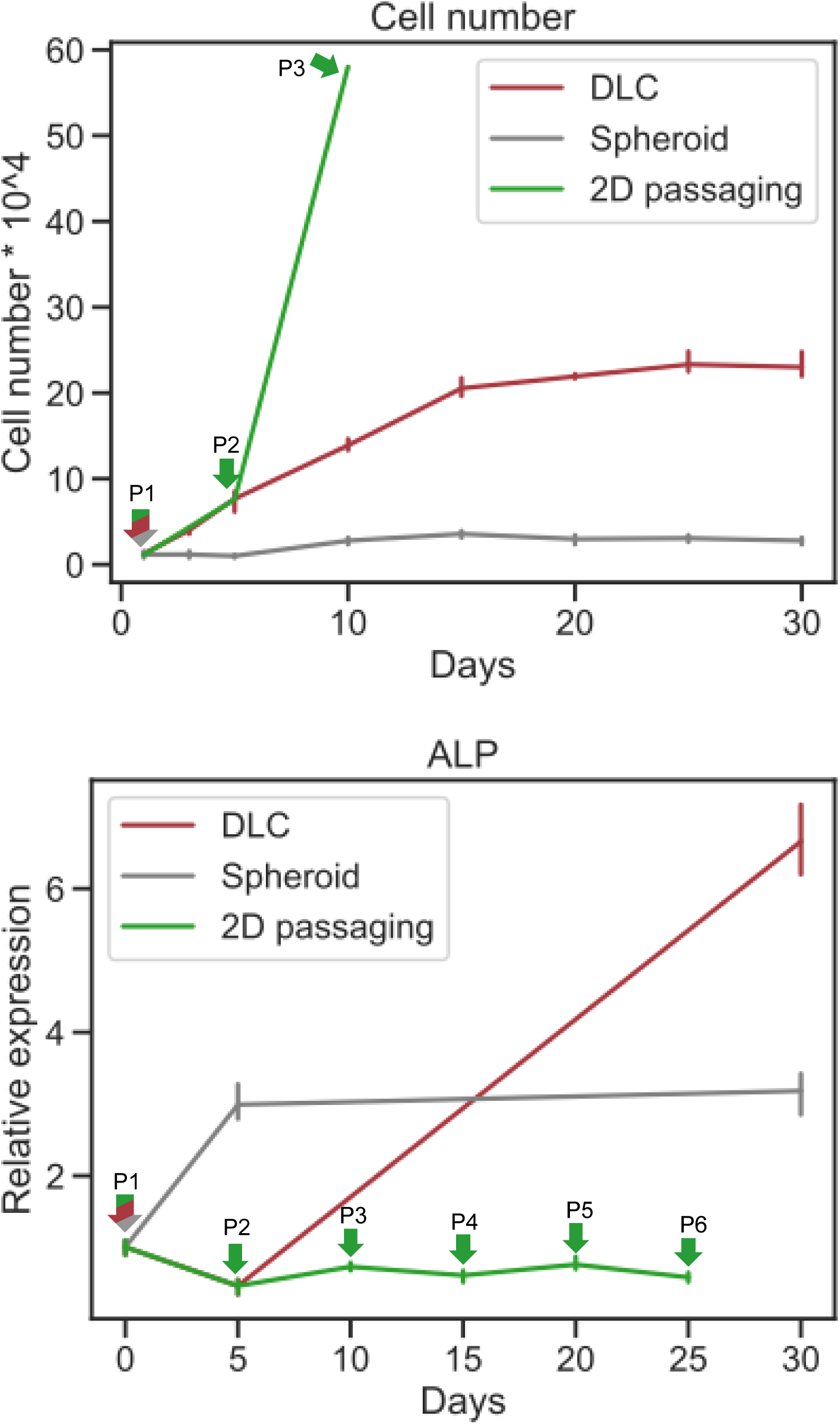
**Comparison of cell proliferation and ALP expression among 2D passaging, spheroid culture, and DLC.** (Top) Growth kinetics of hDPCs cultured under three conditions: 2D monolayer with serial passaging (green), spheroid culture (gray), and dense-layer culture (DLC; red). Cell numbers were quantified over 30 days. Arrows indicate passage points for the 2D condition (P1–P3). (Bottom) Relative ALP gene expression levels were assessed by qPCR at each time point. While 2D-passaged cells showed a decline in ALP expression over successive passages (P1–P6), DLC-cultured hDPCs exhibited a continuous increase, surpassing both 2D and spheroid conditions by day 30.

**Supplementary fig. 2.**
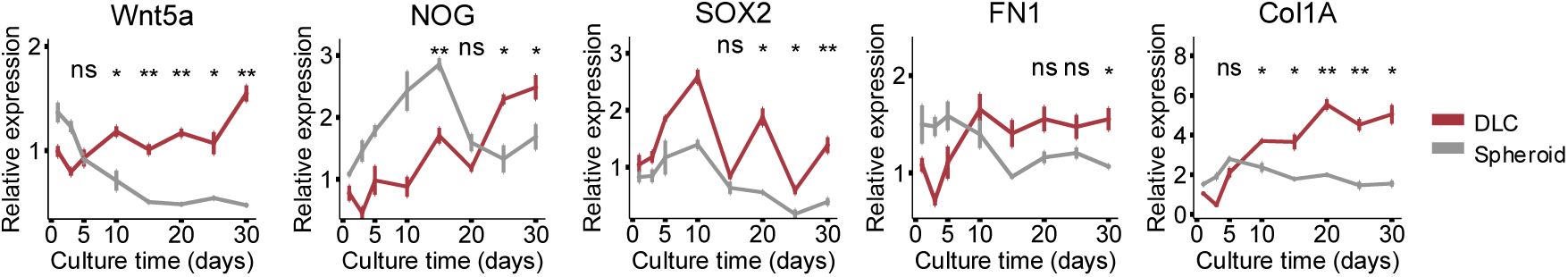
Relative expression of marker genes in DLC- and spheroid-cultured hDPC.

**Supplementary fig. 3.**
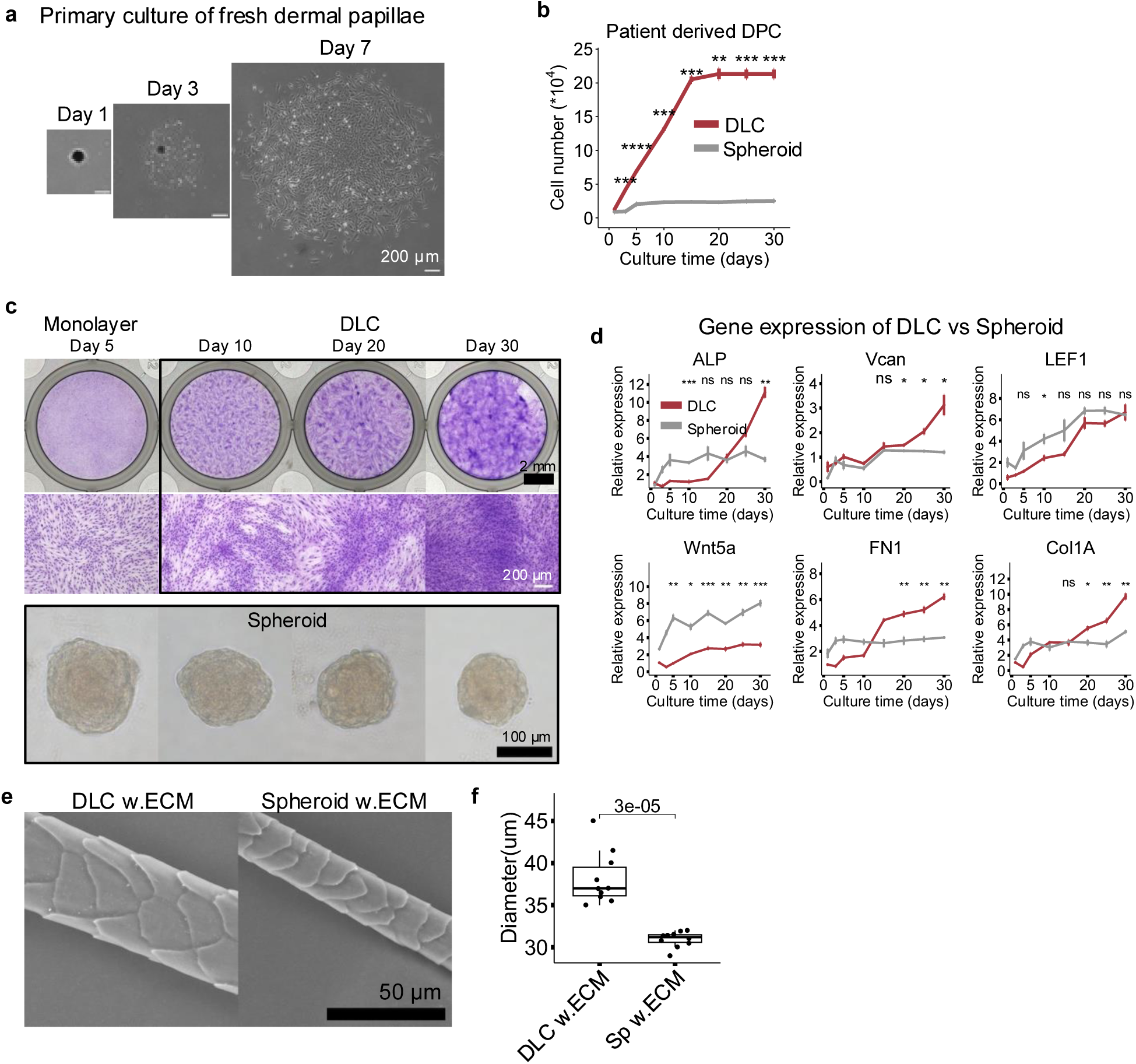
Dense-layered culture of dermal papilla cells isolated from AGA patients. **a** Primary culture of AGA patient-derived dermal papillae **b** Cell proliferation rate of DLC- and spheroid-cultured AGA patientderived hDPC (n = 3) **c** Morphological transformation of DLC-treated and spheroid-cultured AGA patient-derived hDPC **d** Relative expression of genes of DLC- and spheroid-cultured hDPC **e** Cuticle structures of hairs generated by DLC and spheroid cultures. **f** Hair diameters generated by DLC with ECM versus spheroids with ECM. Two-tailed Student’s t-tests (b, d, h). Oneway ANOVA with Tukey’s multiple comparison test. (f) Box plots indicate median (middle line), 25th, 75th percentile (box), and 10th and 90th percentiles (whiskers) as well as outliers.

**Supplementary fig. 4.**
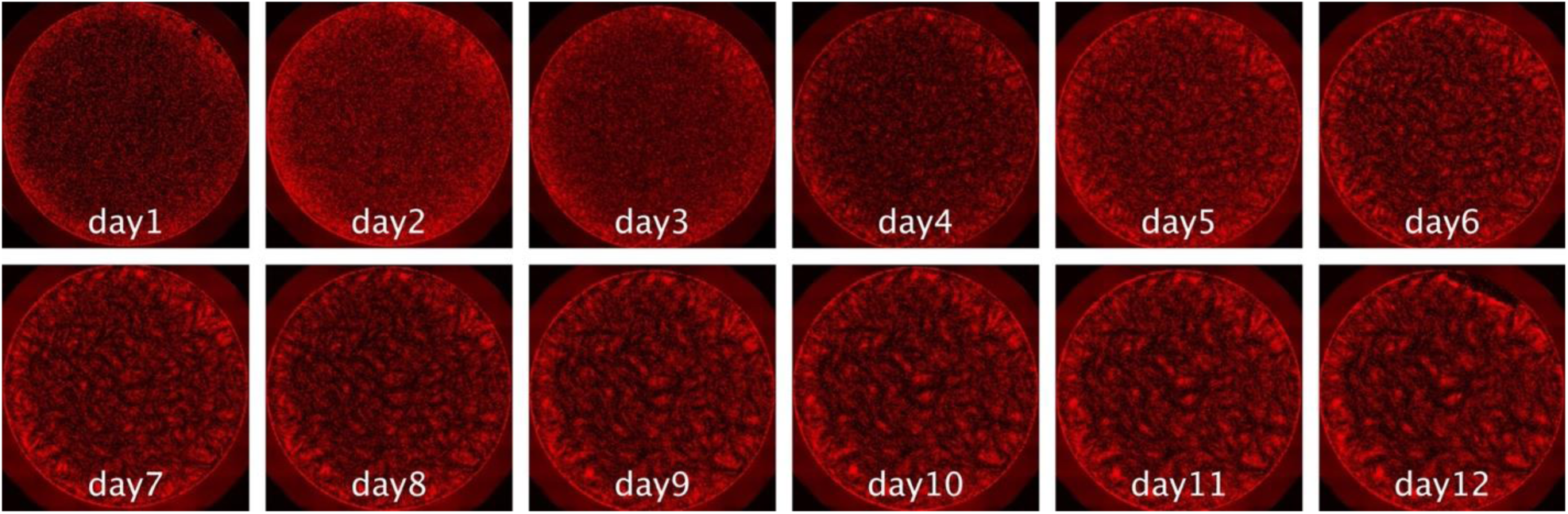
**Live imaging of hDPC for DLC, indicating that ridge area and valley area do not shift drastically after day 6**

**Supplementary fig. 5.**
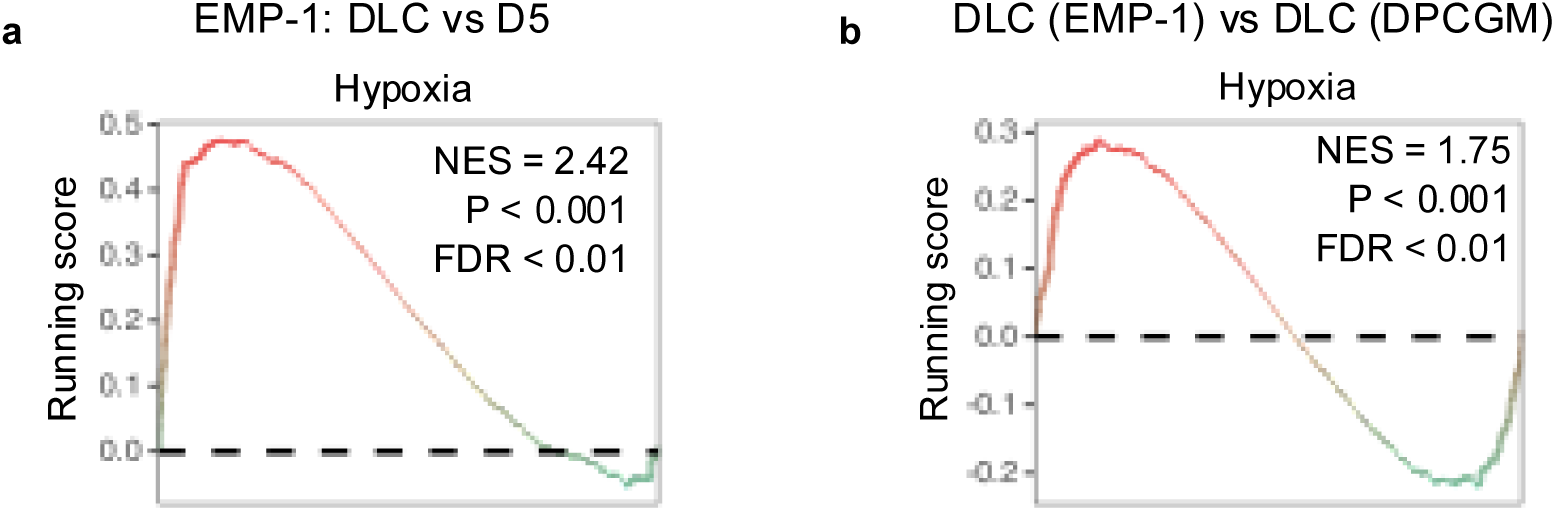
Activation of the hypoxia pathway in hDPCs under different culture medium. **a** Hypoxia signaling pathway activated in hDPCs cultured in EMP-1 medium under DLC condition. **b** EMP-1 medium induces significantly stronger hypoxia pathway activation than DPCGM under DLC condition at day 30 of culture.

**Supplementary Figure 6.**
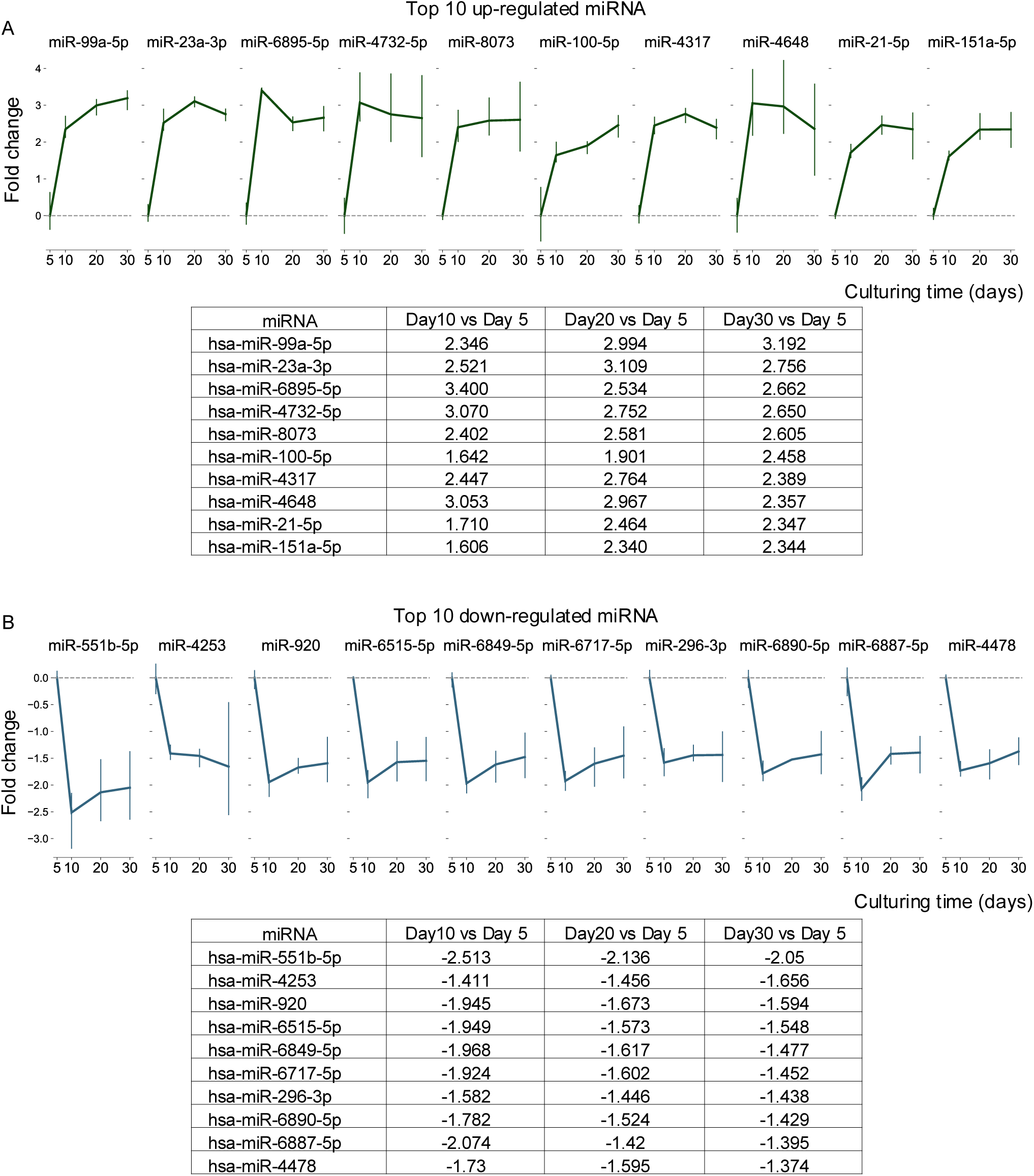

**Supplementary Table 1.** Genes only restored by DLC culture comparing to P0.

|  | Gene<br>Symbol | Fresh<br>logFC | DLC<br>logFC | Spheroid<br>logFC |
| --- | --- | --- | --- | --- |
| 1 | STMN2 | 2.267 | 4.723 | 1.060 |
| 2 | IGF2 | 5.208 | 4.430 | 0.047 |
| 3 | ALPL | 3.525 | 4.396 | 1.219 |
| 4 | RSPO3 | 3.688 | 4.313 | 0.989 |
| 5 | OMD | 5.243 | 4.216 | 1.436 |
| 6 | COL15A1 | 3.691 | 4.036 | 1.640 |
| 7 | SYNPO2 | 2.428 | 3.813 | -0.514 |
| 8 | SVEP1 | 2.159 | 3.710 | 0.796 |
| 9 | CSTA | 3.782 | 3.681 | 1.083 |
| 10 | RNF150 | 1.825 | 3.424 | 0.877 |
| 11 | AFF3 | 2.853 | 3.395 | -0.358 |
| 12 | NEDD9 | 3.155 | 3.289 | -0.157 |
| 13 | DHRS3 | 3.696 | 3.286 | 0.825 |
| 14 | EDIL3 | 1.971 | 3.260 | 1.173 |
| 15 | ADAMTS5 | 2.235 | 3.254 | 0.992 |
| 16 | DSP | 4.931 | 3.250 | -0.267 |
| 17 | C10orf10 | 2.493 | 3.129 | -0.110 |
| 18 | PRUNE2 | 1.606 | 3.070 | -0.334 |
| 19 | MXRA5 | 1.756 | 2.854 | -0.186 |
| 20 | FRY | 3.730 | 2.818 | -0.184 |
| 21 | MAP2K6 | 1.893 | 2.641 | -0.020 |
| 22 | SYTL2 | 2.519 | 2.561 | 0.161 |
| 23 | SYBU | 2.789 | 2.547 | -1.111 |
| 24 | ANK2 | 2.339 | 2.539 | 1.475 |
| 25 | CCL28 | 1.993 | 2.407 | 1.113 |
| 26 | MAF | 1.858 | 2.337 | 1.405 |
| 27 | FMOD | 2.025 | 2.322 | 0.934 |
| 28 | CORIN | 2.499 | 2.313 | 0.353 |
| 29 | PRRG3 | 2.087 | 2.298 | 1.311 |
| 30 | KIT | 3.517 | 2.289 | 0.899 |
| 31 | SOD3 | 2.151 | 2.282 | 0.020 |
| 32 | COBLL1 | 1.886 | 2.152 | -0.028 |
| 33 | LXN | 2.206 | 2.151 | -0.746 |
| 34 | JAG1 | 4.137 | 2.126 | 1.324 |
| 35 | ACVR2A | 1.543 | 2.122 | 0.592 |
| 36 | KCNMB4 | 1.642 | 2.100 | 1.045 |
| 37 | LRIG3 | 1.863 | 2.097 | 0.364 |
| 38 | ATF3 | 4.126 | 2.085 | -0.211 |
| 39 | IGFBP5 | 2.613 | 2.061 | 0.201 |
| 40 | ER | 1.930 | 2.044 | 0.981 |
| 41 | SFRP1 | 3.093 | 2.017 | 0.401 |
| 42 | GPX3 | 3.108 | 1.994 | -0.871 |
| 43 | NUAK1 | 2.321 | 1.979 | 1.427 |
| 44 | SERPING1 | 2.345 | 1.968 | 0.642 |
| 45 | SORT1 | 1.720 | 1.946 | 0.668 |
| 46 | SPATA17 | 1.609 | 1.919 | -1.233 |
| 47 | SSPN | 1.644 | 1.908 | 0.584 |
| 48 | HLA-DMA | 3.700 | 1.896 | -0.251 |
| 49 | CXADR | 5.179 | 1.881 | 0.301 |
| 50 | FAM149A | 1.963 | 1.827 | 1.340 |
| 51 | DMD | 2.418 | 1.824 | -2.102 |
| 52 | NTN4 | 2.200 | 1.824 | -1.189 |
| 53 | PARD3B | 1.791 | 1.808 | -0.225 |
| 54 | SERTAD4 | 3.498 | 1.798 | 1.084 |
| 55 | NIPSNAP3B | 1.795 | 1.778 | 1.248 |
| 56 | MYLIP | 2.284 | 1.776 | 1.379 |
| 57 | LIMA1 | 1.831 | 1.763 | 0.331 |
| 58 | UBE2D4 | 1.744 | 1.734 | 0.844 |
| 59 | LINC00478 | 3.156 | 1.714 | 0.737 |
| 60 | PPL | 3.154 | 1.696 | 0.111 |
| 61 | P2RX7 | 2.784 | 1.687 | 0.917 |
| 62 | FAM198B | 1.850 | 1.684 | 0.947 |
| 63 | MTURN | 1.955 | 1.684 | 1.301 |
| 64 | HSPA4L | 1.616 | 1.676 | 0.657 |
| 65 | SCUBE2 | 4.050 | 1.658 | 0.553 |
| 66 | DIO2 | 4.831 | 1.652 | 0.848 |
| 67 | PAPLN | 3.148 | 1.645 | 0.809 |
| 68 | NTRK2 | 2.719 | 1.640 | 0.554 |
| 69 | SEPP1 | 1.850 | 1.577 | 1.016 |
| 70 | BTG2 | 4.068 | 1.548 | 1.129 |
| 71 | PAWR | 1.711 | 1.525 | -0.651 |
| 72 | PRLR | 2.537 | 1.500 | -0.228 |

**Supplementary Table 2.** Genes only restored by spheroid culture comparing to P0.

|  | Gene<br>Symbol | Fresh<br>logFC | DLC<br>logFC | Spheroid<br>logFC |
| --- | --- | --- | --- | --- |
| 1 | PDK4 | 3.353 | 0.455 | 4.292 |
| 2 | SLC7A8 | 2.711 | 0.468 | 3.897 |
| 3 | PLXDC2 | 2.961 | 1.443 | 3.572 |
| 4 | ID2 | 2.937 | 0.292 | 3.416 |
| 5 | HES1 | 3.988 | 0.685 | 3.341 |
| 6 | FOXO1 | 3.333 | 1.041 | 3.221 |
| 7 | SNCAIP | 3.851 | 0.744 | 3.174 |
| 8 | BMP2 | 2.717 | -0.606 | 3.097 |
| 9 | ID3 | 3.861 | -0.641 | 3.066 |
| 10 | SOX4 | 2.055 | 1.310 | 3.063 |
| 11 | RGS2 | 2.257 | -0.665 | 2.891 |
| 12 | SERPINI1 | 1.692 | -1.016 | 2.859 |
| 13 | BAMBI | 4.670 | 1.480 | 2.836 |
| 14 | TGFB3 | 2.587 | 1.481 | 2.673 |
| 15 | KDM7A | 2.115 | 0.665 | 2.544 |
| 16 | UNC5B | 2.562 | 1.175 | 2.468 |
| 17 | VASH2 | 1.838 | 0.430 | 2.466 |
| 18 | MATN2 | 3.408 | 1.371 | 2.433 |
| 19 | TFAP2A | 3.351 | 1.024 | 2.432 |
| 20 | FNIP2 | 1.698 | 1.480 | 2.404 |
| 21 | SMAD9 | 2.618 | -0.482 | 2.347 |
| 22 | EFCAB6 | 1.806 | 0.485 | 2.346 |
| 23 | MTUS1 | 2.485 | 0.908 | 2.330 |
| 24 | PMEPA1 | 1.966 | 0.128 | 2.286 |
| 25 | SERINC5 | 1.779 | 1.361 | 2.285 |
| 26 | CLCA2 | 4.847 | 1.341 | 2.226 |
| 27 | PDK3 | 2.101 | 1.069 | 2.179 |
| 28 | HEY1 | 4.579 | -0.379 | 2.174 |
| 29 | MXD1 | 1.607 | 1.041 | 2.159 |
| 30 | PABPC4L | 2.029 | 1.078 | 2.132 |
| 31 | BCL2L11 | 1.814 | 0.763 | 2.117 |
| 32 | FMN1 | 1.946 | 0.673 | 2.106 |
| 33 | JHDM1D-AS1 | 1.690 | 0.956 | 2.075 |
| 34 | HSPB8 | 3.803 | 1.108 | 2.050 |
| 35 | PPM1L | 1.597 | 1.382 | 2.036 |
| 36 | KLHL41 | 3.340 | 1.174 | 2.013 |
| 37 | PLK2 | 2.441 | 1.379 | 1.950 |
| 38 | SAMD5 | 2.731 | 1.183 | 1.908 |
| 39 | CALCRL | 3.508 | -0.900 | 1.852 |
| 40 | SETBP1 | 1.876 | 1.034 | 1.840 |
| 41 | CDKN1C | 1.879 | 1.168 | 1.823 |
| 42 | TXNDC16 | 1.593 | 1.021 | 1.821 |
| 43 | TGFB2 | 2.090 | 1.287 | 1.800 |
| 44 | CADM1 | 2.831 | 1.498 | 1.767 |
| 45 | MITF | 3.346 | 1.467 | 1.757 |
| 46 | DOK6 | 1.554 | 1.464 | 1.716 |
| 47 | VAV3 | 2.464 | -0.545 | 1.713 |
| 48 | KAZN | 1.837 | 0.978 | 1.683 |
| 49 | SMAD6 | 2.205 | 0.475 | 1.653 |
| 50 | LOC728392 | 2.085 | 1.298 | 1.648 |
| 51 | LOC102723845 | 2.476 | 0.863 | 1.631 |
| 52 | SNN | 1.748 | 1.412 | 1.619 |
| 53 | BGN | 1.931 | 0.854 | 1.605 |
| 54 | NPL | 1.997 | 0.395 | 1.605 |
| 55 | TMEM133 | 1.512 | 0.996 | 1.604 |
| 56 | FAM213A | 1.507 | 0.840 | 1.588 |
| 57 | PTH1R | 3.062 | 0.813 | 1.560 |
| 58 | SKIL | 1.891 | 0.955 | 1.516 |
| 59 | GAS7 | 1.812 | 0.675 | 1.514 |

**Supplementary Table 3.** Genes restored by both DLC culture and spheroid culture.

|  | Gene Symbol | logFC | logFC | logFC | Comparison |
| --- | --- | --- | --- | --- | --- |
| 1 | ECM2 | 3.167 | 4.229 | 1.870 | DLC>Spheroid |
| 2 | MAN1C1 | 2.259 | 3.959 | 1.638 | DLC>Spheroid |
| 3 | ST6GALNAC5 | 3.711 | 5.116 | 2.969 | DLC>Spheroid |
| 4 | COL8A2 | 3.154 | 4.210 | 2.267 | DLC>Spheroid |
| 5 | ASPN | 4.638 | 3.954 | 2.035 | DLC>Spheroid |
| 6 | PDGFD | 2.388 | 3.245 | 1.780 | DLC>Spheroid |
| 7 | FGFR2 | 5.039 | 3.539 | 2.301 | DLC>Spheroid |
| 8 | BHLHE41 | 4.665 | 3.079 | 1.862 | DLC>Spheroid |
| 9 | THRB | 2.766 | 3.705 | 2.607 | DLC>Spheroid |
| 10 | LRRC15 | 3.375 | 2.856 | 1.799 | DLC>Spheroid |
| 11 | ANK2 | 2.822 | 2.579 | 1.593 | DLC>Spheroid |
| 12 | SULF1 | 2.717 | 2.887 | 1.993 | DLC>Spheroid |
| 13 | STON2 | 3.517 | 3.660 | 2.925 | DLC>Spheroid |
| 14 | COL14A1 | 2.862 | 3.618 | 2.948 | DLC>Spheroid |
| 15 | MFRP | 2.756 | 2.867 | 2.231 | DLC>Spheroid |
| 16 | FAM134B | 2.473 | 2.968 | 2.385 | DLC>Spheroid |
| 17 | NUPR1 | 2.334 | 2.446 | 1.962 | DLC>Spheroid |
| 18 | RDH10 | 2.117 | 2.603 | 2.163 | DLC>Spheroid |
| 19 | DAPK1 | 3.154 | 2.497 | 2.075 | DLC>Spheroid |
| 20 | EXPH5 | 2.828 | 3.222 | 2.870 | DLC>Spheroid |
| 21 | IGFBP3 | 3.500 | 3.114 | 2.982 | DLC>Spheroid |
| 22 | PCDHB6 | 1.732 | 1.899 | 1.820 | DLC>Spheroid |
| 23 | KIF26B | 3.563 | 3.485 | 3.429 | DLC>Spheroid |
| 24 | BCL2 | 3.026 | 1.595 | 1.581 | DLC>Spheroid |
| 25 | SULF2 | 2.981 | 2.816 | 2.807 | DLC>Spheroid |
| 26 | ENAH | 1.909 | 2.206 | 2.297 | Spheroid>DLC |
| 27 | BHLHE40 | 3.418 | 2.346 | 2.565 | Spheroid>DLC |
| 28 | ZIC1 | 2.864 | 1.521 | 1.743 | Spheroid>DLC |
| 29 | DPT | 3.165 | 2.291 | 2.545 | Spheroid>DLC |
| 30 | OLFM2 | 1.616 | 2.475 | 2.770 | Spheroid>DLC |
| 31 | PDE1A | 2.974 | 2.855 | 3.159 | Spheroid>DLC |
| 32 | CPE | 2.144 | 2.111 | 2.422 | Spheroid>DLC |
| 33 | MITF | 2.928 | 1.569 | 1.956 | Spheroid>DLC |
| 34 | MTSS1 | 3.504 | 1.551 | 2.007 | Spheroid>DLC |
| 35 | PCDH7 | 2.084 | 3.332 | 3.810 | Spheroid>DLC |
| 36 | SLC1A4 | 2.882 | 2.236 | 2.741 | Spheroid>DLC |
| 37 | PCDHB9 | 2.068 | 1.823 | 2.356 | Spheroid>DLC |
| 38 | FRZB | 6.737 | 2.003 | 2.543 | Spheroid>DLC |
| 39 | TGFB2 | 3.071 | 1.891 | 2.596 | Spheroid>DLC |
| 40 | PPM1L | 1.518 | 1.589 | 2.303 | Spheroid>DLC |
| 41 | ZNF704 | 3.708 | 1.662 | 2.411 | Spheroid>DLC |
| 42 | VLDLR | 2.332 | 2.683 | 3.512 | Spheroid>DLC |
| 43 | SLC40A1 | 4.515 | 3.488 | 4.400 | Spheroid>DLC |
| 44 | NTRK2 | 6.257 | 2.207 | 3.541 | Spheroid>DLC |
| 45 | ID4 | 5.512 | 2.407 | 3.986 | Spheroid>DLC |
| 46 | GPC4 | 4.172 | 1.656 | 3.316 | Spheroid>DLC |
| 47 | MMP11 | 1.937 | 2.293 | 4.486 | Spheroid>DLC |
| 48 | PLXDC2 | 2.890 | 1.570 | 4.154 | Spheroid>DLC |

**Supplementary Table 4.**
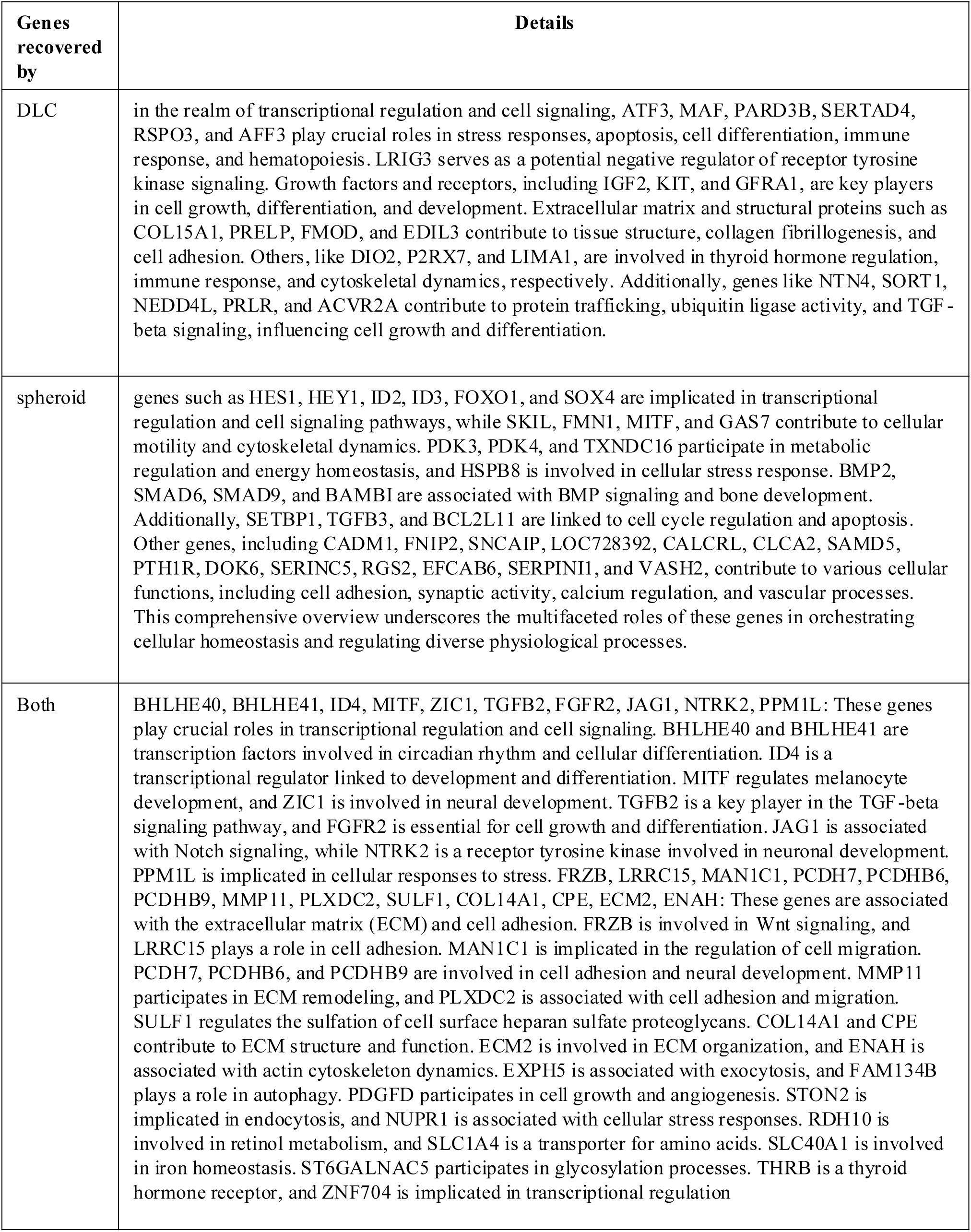
Details of genes restored by each DLC, spheroid culture and both.

